# Using subthreshold events to characterize the functional architecture of electrically coupled networks

**DOI:** 10.1101/483156

**Authors:** Yaara Lefler, Oren Amsalem, Idan Segev, Yosef Yarom

## Abstract

The electrical connectivity in the inferior olive (IO) nucleus plays an important role in generating well-timed spiking activity. Here we combined electrophysiological and computational approaches to assess the functional organization of mice IO nucleus. Spontaneous fast and slow subthreshold events were commonly encountered during in vitro recordings. We show that the fast events represent a regenerative response in unique excitable spine-like structures in the axon hillock, whereas the slow events reflect the electrical connectivity between neurons (‘spikelets’). Recordings from cell pairs revealed the synchronized occurrence of distinct groups of spikelets; their rate and distribution enabled an accurate estimation of the number of connected cells and is suggestive of a clustered organization. This study thus provides a new perspective on the functional and structural organization of the olivary nucleus, insights into two different subthreshold non-synaptic events, and a novel experimental and theoretical approach to the study of electrically-coupled networks.

## Introduction

In recent years research has confirmed that electrically coupled neural networks are found in every major region of the central nervous system (Condorelli *et al.*, 2000; M. V.. Bennett and Zukin, 2004; Connors and Long, 2004; Hormuzdi *et al.*, 2004). One common feature of these networks is their synchronized rhythmic activity (Connors and Long, 2004; M. V. L. Bennett and Zukin, 2004; Connors, 2017; Coulon and Landisman, 2017) which has been shown to be correlated with higher brain functions such as states of arousal, awareness, cognition, and attention (Ritz and Sejnowski, 1997; Engel, Fries and Singer, 2001; Buzsáki, 2005; Steriade, 2006; Uhlhaas *et al.*, 2009; Wang, 2010). Recently it has been demonstrated that the efficiency of electrical synapses is modulated by electrical and chemical activity, very much like chemical synapses (O’brien, 2014; Marder, Gutierrez and Nusbaum, 2016; Coulon and Landisman, 2017). It thus stands to reason that the functional architecture of these networks must undergo continuous modification to meet system demands. This underscores the urgent need to determine the functional state of a network and associate it with the corresponding brain states. Since anatomical information is insufficient, this can only be done using physiological parameters that capture the functional architecture of a network at any given time.

The inferior olive electrically coupled network, which was among the first networks to be studied in the mammalian brain, provides primary excitatory input to the cerebellar cortex (Eccles, Llinás and Sasaki, 1966). There is a general consensus that the function of this network is to generate synchronous activity of the olivary neurons, which provide temporal information for either learning processes, motor execution, sensory predictions or expectations (Llinas and Sasaki, 1989; Lou and Bloedel, 1992; Welsh *et al.*, 1995; Van Der Giessen *et al.*, 2008; Llinás, 2009; De Zeeuw *et al.*, 2011). Temporal information is thought to be generated by the subthreshold sinusoidal-like oscillations of the membrane voltage that appear to emerge from an interplay between the membrane properties and network connectivity (Llinas and Yarom, 1986; Lampl and Yarom, 1997; Manor *et al.*, 1997; Loewenstein, Yarom and Sompolinsky, 2001; Devor and Yarom, 2002b). Recently this oscillatory activity was shown to be governed by synaptic inputs that partially originate in the deep cerebellar nuclei, and modulate the efficacy of the coupling, by defining the spatial extent of the electrically coupled network (Lefler, Yarom and Uusisaari, 2014; Mathy, Clark and Häusser, 2014; Turecek *et al.*, 2014).

Early work on the morphological organization of the IO indicated that it is organized in clusters of up to 8 neurons, whose dendrites are integrated in glomerulus structures (Sotelo, Llinas and Baker, 1974) and are innervated by both excitatory and inhibitory synaptic inputs (De Zeeuw *et al.*, 1990). This presumed cluster organization has been supported by dye coupling studies showing that each olivary neuron is anatomically coupled to roughly ten other neurons (Devor and Yarom, 2002a; Leznik and Llinas, 2005; Placantonakis *et al.*, 2006; Hoge *et al.*, 2011; Turecek *et al.*, 2014). However, the organization of the network has only been addressed physiologically in a few voltage-sensitive dye imaging studies which found ensembles of synchronously active neurons corresponding to a cluster size estimation of hundreds of neurons (Devor and Yarom, 2002b; Leznik, Makarenko and Llinas, 2002). The documented synchronicity of complex spikes activity in tens of cerebellar Purkinje cells during motor tasks and sensory stimulation, is also in favor of such of ensemble organization (Bloedel and Ebner, 1984; Welsh *et al.*, 1995; Welsh and Llinás, 1997; Mukamel, Nimmerjahn and Schnitzer, 2009; Ozden *et al.*, 2009; Schultz *et al.*, 2009; De Zeeuw *et al.*, 2011).

In this study, we describe a novel method to estimate the size and efficacy of a network by analyzing the all-or-none subthreshold unitary activity known as a ‘spikelet’. Initially, spikelets were considered as the manifestation of an action potential transmitted via electrical synapses (Llinas, Baker and Sotelo, 1974; MacVicar and Dudek, 1981; Valiante *et al.*, 1995; Galarreta and Hestrin, 1999; Gibson, Belerlein and Connors, 1999; Mann-Metzer and Yarom, 1999; Hughes *et al.*, 2002; Chorev and Brecht, 2012). However, in other studies, spikelets were referred to either local dendritic regenerative response (Spencer and Kandel, 1961; Golding and Spruston, 1998; Smith *et al.*, 2013), a reflection of an action potential in the initial segment or at an ectopic site along the axon that fails to invade the soma (Stasheff, Hines and Wilson, 1993; Avoli, Methot and Kawasaki, 1998; Juszczak and Swiergiel, 2009; Sheffield *et al.*, 2011; Dugladze *et al.*, 2012; Michalikova, Remme and Kempter, 2017), an electrical coupling between axons (Schmitz *et al.*, 2001; Traub *et al.*, 2002) or simply extracellularly recorded activity of nearby neurons (Vigmond *et al.*, 1997; Scholl *et al.*, 2015). Here we show that the spontaneous unitary events recorded from olivary neurons can be classified into two groups that differ in their waveform and properties: fast events having identical waveforms with variable high amplitudes, and slow events having different waveforms and low amplitudes. We show that the low amplitude slow events reflect the occurrence of action potentials in electrically coupled neurons, whereas the high amplitude fast events are regenerative responses that are likely to represent action potentials that occur at the axonal spines. We then used the slow events that were simultaneously recorded in pairs of neurons to estimate the number of neurons in the network. We found that each olivary neuron is electrically connected to an average of 19 other neurons and that the network is not randomly connected, rather it is composed of functional clusters of connected neurons.

## Results

### Spontaneous unitary events recorded in neurons of the inferior olive

The subthreshold spontaneous activity recorded from IO neurons (Figure 1A) was characterized by unitary unipolar events that varied in amplitude and waveform. This spontaneous activity, which was observed in 74.3% of the neurons (188 out of 253) with an average rate of 0.7±0.6/sec (calculated in 70 neurons) were also encountered in the presence of excitatory synaptic blockers (CNQX and/or APV, n=19) although the frequency of occurrence was reduced to an average below 0.02 Hz. These subthreshold events could readily be divided into two populations of small and large events (Figure 1A inset, circles vs. stars), as shown by the amplitude histogram (Figure 1B). K-means clustering of the event waveforms reveals 5 distinct groups (Figure 1C), which when normalized (Figure 1D) showed the waveform difference between the two types; one type had high amplitude and fast kinetics, and the second type had low amplitude and slow kinetics.

**Figure 1.**
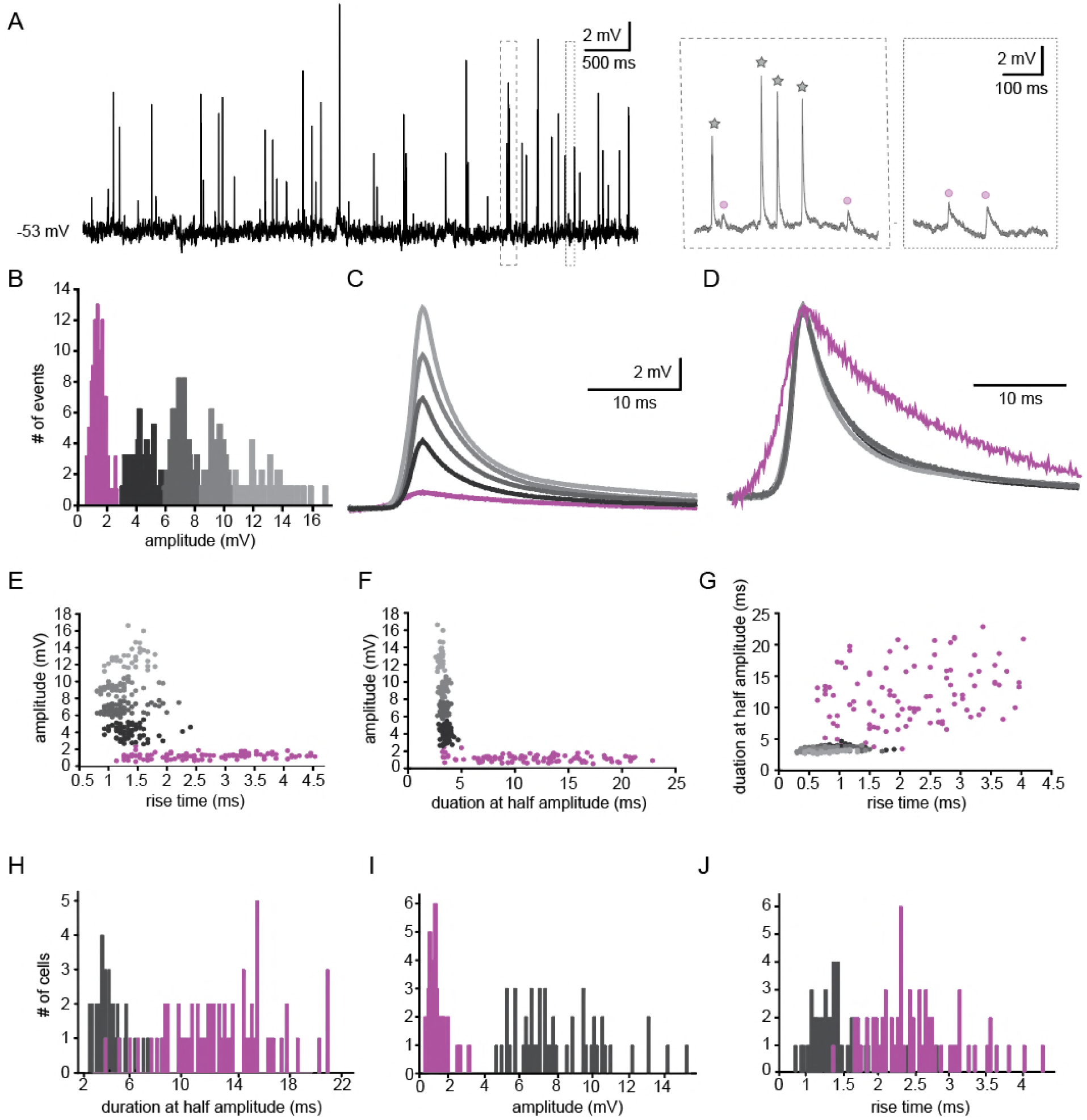
Two types of subthreshold spontaneous events recorded in olivary neurons. (A) Spontaneous subthreshold events recorded from an olivary neuron. Right panels - higher magnification of the marked rectangles; black stars - fast and high amplitude events; purple circles - slow and small events. (B) the distribution of the event amplitudes; colors were assigned according to the K-means analysis of the amplitudes. (C) Averages of the subthreshold events in each cluster, color coded as in B. (D) the normalized events shown in c. (E-G) Scatter plots for the relationships between the shape indices of the subthreshold events (color coded as in B). (E) Amplitude and rise time; (F) Amplitude and half width; (G) half width and rise time. (H-J) Histograms of the shape indices (half width (H); amplitude (I); and rise time (J)) of the subthreshold events in a population of 49 olivary neurons; purple and gray bars correspond to slow and fast events respectively.

To further analyze the event waveforms, we measured each event’s rise time and duration at half amplitude. The results obtained from a representative neuron are summarized in Figure 1 E-G. One type (black to grey circles) had a relatively high amplitude (2.4-16.6 mV; average of 7.5±3.1 mV) and fast kinetics (average rise time of 1.3±0.3 ms and average half duration of 3.4±0.3 ms) whereas the second type (purple circles) had a relatively low amplitude (<2mV: 0.6-1.9 mV; average of 1.17±0.3 mV) and slow kinetics (average rise time of 2.4±0.4 ms and average half duration of 11.8±5 ms). Plotting the duration as a function of the rise time (Figure 1G), which further supports the two-type scheme, failed to demonstrate a monotonous relationship between the dendritic location of the synapse and the rise-time/half-width expected from Rall’s cable theory (Rall, 1967). Thus, it seems unlikely that the two types represent signals arising from different locations along the cell’s morphological structure. The distribution of rise-time and half-width in a population of 49 neurons, which is summarized in Figure 1H-J, confirms that there were indeed two distinct types of events. Whereas the high amplitudes events had a fast rise time (0.8-2.8 ms; average of 1.4±0.4 ms) and short duration (2.5-8.3 ms; average of 4.2±1.3 ms), the low amplitude events had a longer rise time (1.3-4.3 ms; average of 2.5±0.6 ms) and a longer duration (3.6-21 ms; average of 12.7±3.9 ms). For the high-amplitude events, the broad distribution of amplitudes (ranging from 4.5 to15.3 mV) and the somewhat limited distribution of rise-times and durations strongly indicates that these groups of fast events were generated by a similar mechanism.

Overall, the frequency of slow events was four times higher (0.56±0.62 /sec; n=69) than that of the fast events (0.14±0.18 /sec; n=58). It should be noted however, that due to low amplitude and limited resolution, a further division of the slower type was difficult However, in 9 experiments, this slow signal could be subdivided into two groups showing similar slow kinetics with different amplitudes (not shown).

### Slow events reflect electrical coupling between neurons

Whole-cell recordings from pairs of coupled olivary neurons revealed that the post synaptic responses to either spontaneously (Figure 2A) or evoked (Figure 2B) action potentials in one neuron were precisely correlated with depolarizing events in the coupled neuron. Both the spontaneous and the evoked events resembled the spontaneously recorded slow events depicted in Figure 1. These events had an amplitude of 1.2±0.12 mV, a rise time of 3.7±0.9 ms and a duration of 14.9±4.0 ms, thus well within the range of spontaneously measured slow events. Paired recordings from 36 neurons showed that the amplitudes of the events varied from 0.2 to1.67 mV (average of 0.88±0.51 mV) whereas the average rise times and half durations were 2.95±1.05 ms and 21.43±11.05 ms, respectively, in line with the measured distribution of spontaneously slow events (Figure 1H-J). Importantly, spike-triggered depolarizing events for each pair showed minimal variations. Therefore, the relatively large variability of the spontaneous slow events (Figure 1E-G) is likely to represent action potentials in many neurons that were coupled to the recorded neuron. It should be noted that in these paired recordings we never encountered an evoked postsynaptic response that resembled the fast events depicted in Figure 1.

**Figure 2.**
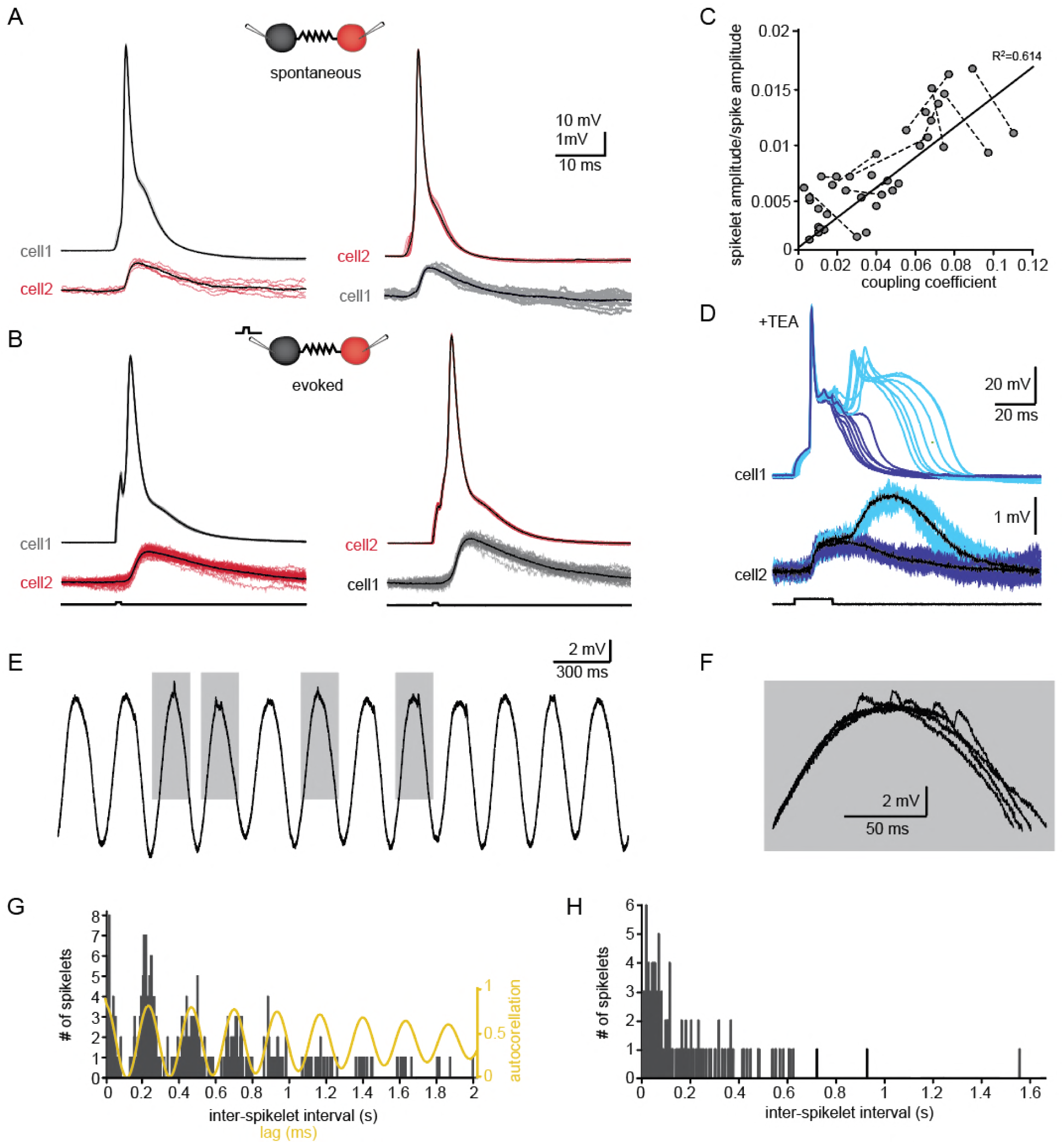
The slow events represent the electrical coupling between neurons. (A) superimposed traces of spontaneously occurring action potentials recorded simultaneously from a pair of coupled neurons (red and gray traces; black traces represent the average events). (B) the same as in A for action potentials evoked by 100 pA, 1 ms current pulses. (C) The linear relationship between the DC coupling coefficient and the spike coupling coefficient. Pairs of cells are connected by dashed lines. Blue line is the linear regression fit (R^2^=0.6). (D) Paired recording in the presence of 10mM TEA. Action potentials with relatively long durations (upper panel, blue traces) were elicited in cell 1 by 50 pA,20 ms current pulse. Occasionally they were followed by a second response (cyan traces). These action potentials elicited post junctional responses in cell 2 with corresponding waveforms (lower traces). (E) Subthreshold events recorded in oscillating olivary neuron. (F) Superposition of the gray rectangles in E, at higher magnification. Note that spikelets were only present for 50 ms along the peak of the oscillations. (G) Inter-spikelet interval (ISLI) from the same neuron (using 4 ms bins), and an autocorrelation (yellow line) of the membrane potential. (H) The ISLI distribution in a non-oscillating neuron.

The relatively broad range of spikelet parameters (Figure 1) can be attributed to a wide range of coupling strengths, different locations of the gap-junctions along the dendritic structure or different durations and shapes of the pre-junctional action potential which is a well-known feature of olivary action potentials (Llinás and Yarom, 1981a, 1981b). We first examined the effect of coupling strength by calculating the ratio of the amplitudes of the pre-junctional action potential to the post-junctional spikelet, and compared it to the coupling coefficient measured by direct current injection (see Methods). As shown in Figure 2C, there was a significant positive correlation (with a slope of 0.134; R2=0.614, p<0.0001; Pearson correlation). Next, we examined the effect of the shape of the pre-junctional action potential on the spikelet parameters. To that end, we partially blocked the voltage dependent potassium current by adding TEA (10mM) to the bath solution. In the presence of TEA, a variety of action potential waveforms were elicited by current injection (Figure 2D). In particular, the initial upstroke of the action potential was unaffected, but there was a significant broadening of the repolarizing phase (Figure 2D, upper panel, dark blue) that often elicited a second calcium-dependent action potential (Figure 2D, upper panel, light blue). This variety of action potential waveforms were always associated with postsynaptic responses that could be clustered into two distinct groups (Figure 2D, lower panel). The prolongation of the action potential was, as expected, followed by a matching increase in the duration of the post-junctional responses (Figure 2D, lower panel, dark blue traces). The appearance of the second component was associated with a slow wave of depolarization in the post-junctional cell (light blue traces). This suggests that the wide range of spikelet parameters (Figure 1) can be accounted for by the variability in coupling strength and pre-junctional action potential waveforms.

Finally, we examined the occurrence of spikelets in neurons that exhibited subthreshold oscillatory activity. Since these oscillations occurred simultaneously in several neurons (Lefler, Torben-Nielsen and Yarom, 2013), it was expected that spikelet occurrence will be correlated with the oscillatory activity. About 50% of the olivary neurons showed spontaneous subthreshold oscillations (Figure 2E). Careful examination of the peaks of the oscillations (Figure 2F) revealed that they were crowned with spikelets. To quantify this observation, we calculated the distribution of the inter-spikelet-interval (ISLI, Figure 2G, black bars), and found distinct groups appearing at intervals of 200 ms. We then calculated the autocorrelation function of the subthreshold oscillations (Figure 2G, yellow line) and found that it matched the ISLI perfectly. It is important to note that a similar fit was observed in 60% of the oscillating neurons (n=18) whereas in non-oscillating neurons (n=70) the ISLI exhibited a Poisson-like distribution (Figure 2H). The strong correlation between oscillatory behavior and the occurrence of spikelets further supports the conclusion that these events represent activity in adjacent electrically coupled neurons.

To summarize, the observations in Figures 1 and 2 strongly suggest that the small and slow spontaneous events represent action potentials occurring in electrically coupled neurons and therefore we will refer to them as ‘spikelets’.

### The large and fast events reflect localized regenerative responses

The source of the fast events is a mystery. On the one hand, they seem to be independent of chemical synaptic transmission, and on the other, they are not activated by pre-junctional action potentials in coupled neurons. To resolve this mystery, we first examined the effect of the membrane voltage on the occurrence and waveform of both types of unitary events. We next used an optogenetic approach combined with pharmacological manipulations to gain insights into the source of these events. Finally, we used a computational approach to examine possible mechanisms to account for the experimental observations.

The effect of membrane potential was examined by DC current injection, which on average set the membrane potential to a range of -33 to -90mV. Figure 3A-B shows the aligned superimposed traces of spikelets (Figure 3A) and fast events (Figure 3B) from one neuron. Normalizing the event amplitudes (Figure 3A-B, right panels) shows that whereas the spikelet shape was unaffected by the current injection (A), the fast events showed a slowdown of the late repolarizing phase with hyperpolarization (B). Quantifying the effect of the injected current (see Methods) on the amplitude (Figure 3C-D) and duration at 20% of the amplitude (Figure 3E-F) in 16 neurons revealed no effect on the amplitude of either type of events (average slope R^2^ = 0.0057 and 0.0035, respectively). The spikelet duration was slightly, but not significantly, affected (Figure 3E; R^2^ = 0.44; one-sample t-test p=0.057). In contrast, it significantly increased the duration of the fast events (Figure 3F; R^2^ = 0.712; one-sample t-test p=0.0005). Comparing the two sets of data revealed a significant difference (Figure 3E vs. F; paired t-test p=0.017). Finally, we measured the effect of the DC current injection on the rate of occurrence of the subthreshold unitary events (Figure 3G-H). Whereas the frequency of the spikelets remained unaffected (Figure 3G, R^2^ = 0.012), the frequency of the fast events increased by a factor of up to 5 (Figure 3H; R^2^ = 0.836). This difference between the occurrence of spikelets and the fast events was highly significant (Figure 3G vs. H, p=0.005, paired t-test). These differences further support our presumption that two different mechanisms generate the two types of subthreshold unitary events. Since spikelets are evoked by action potentials in paired cells, DC current injection to the post-junctional cell will not have a significant effect on the number of action potentials in the pre-junctional cell. On the other hand, the voltage dependency of the fast events strongly suggests that they reflect regenerative responses initiated within the neuron. We therefore refer to them as Internal Regenerative Events or IREs.

**Figure 3.**
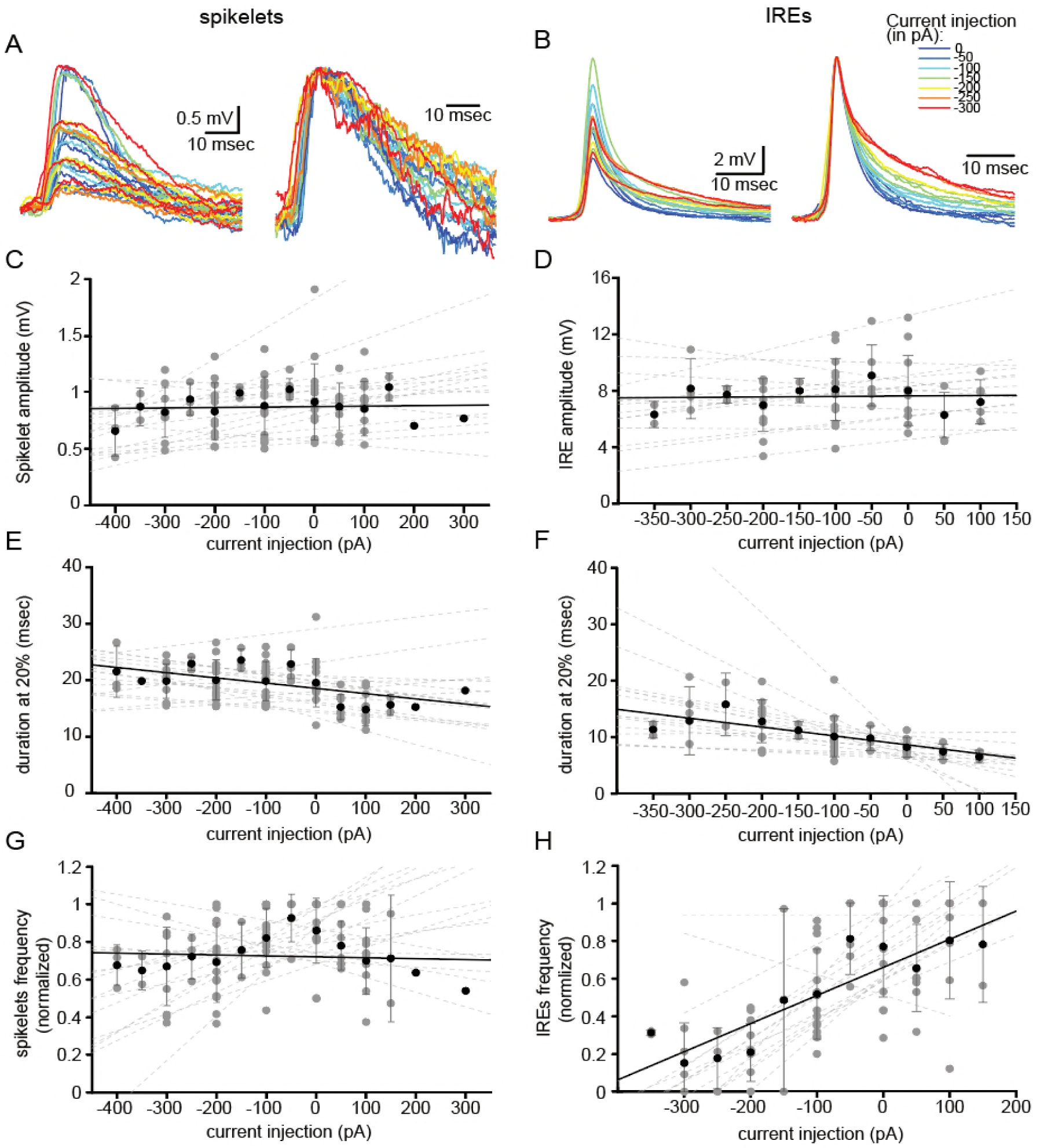
Voltage dependency of the subthreshold events. (A) superimposed spikelets recorded during seven different DC current injections (0 to -300 pA, color coded, left panel), and normalized by amplitude (right panel). (B) same as a, for the fast events (‘IREs’). (C-H) The effect of DC current injection on the amplitude (C-D), the duration, measured at 20% of the amplitude, (E-F) and the frequency of occurrence (G-H) measured in 16 neurons. Gray circles represent the average data from individual neurons, each fitted with a linear regression (dashed gray lines). Black circles and error bars (std) represent the average value for all the neurons in each current injection. Note that the decrease in duration (F) and the increase in frequency (H) with depolarization only occurs for IREs.

Next, we examined the ability of the IREs to trigger a full-blown action potential. Figure 4A shows the superposition of spontaneously occurring action potentials (black and purple traces) and IREs (gray traces) measured from the same neuron. It is clear that some of the action potentials (purple) are preceded by a fast pre-potential that resembles the spontaneous IREs. This possibility was further supported by phase plotting the action potential (Figure 4B, purple and black traces), which showed that 62% of the spontaneous action potentials were triggered by depolarizing events that resembled the IREs in amplitude and waveform. On the population level 29.2±26.7% of the IREs triggered action potentials, whereas 41.6±21.2% of the action potentials were triggered by IREs (Figure 4C). It should be noted that this is an underestimation, since action potentials that seem to arise from the baseline might also be evoked by an IRE that is masked by the beginning of the action potential (see inset in Figure 4A).

**Figure 4.**
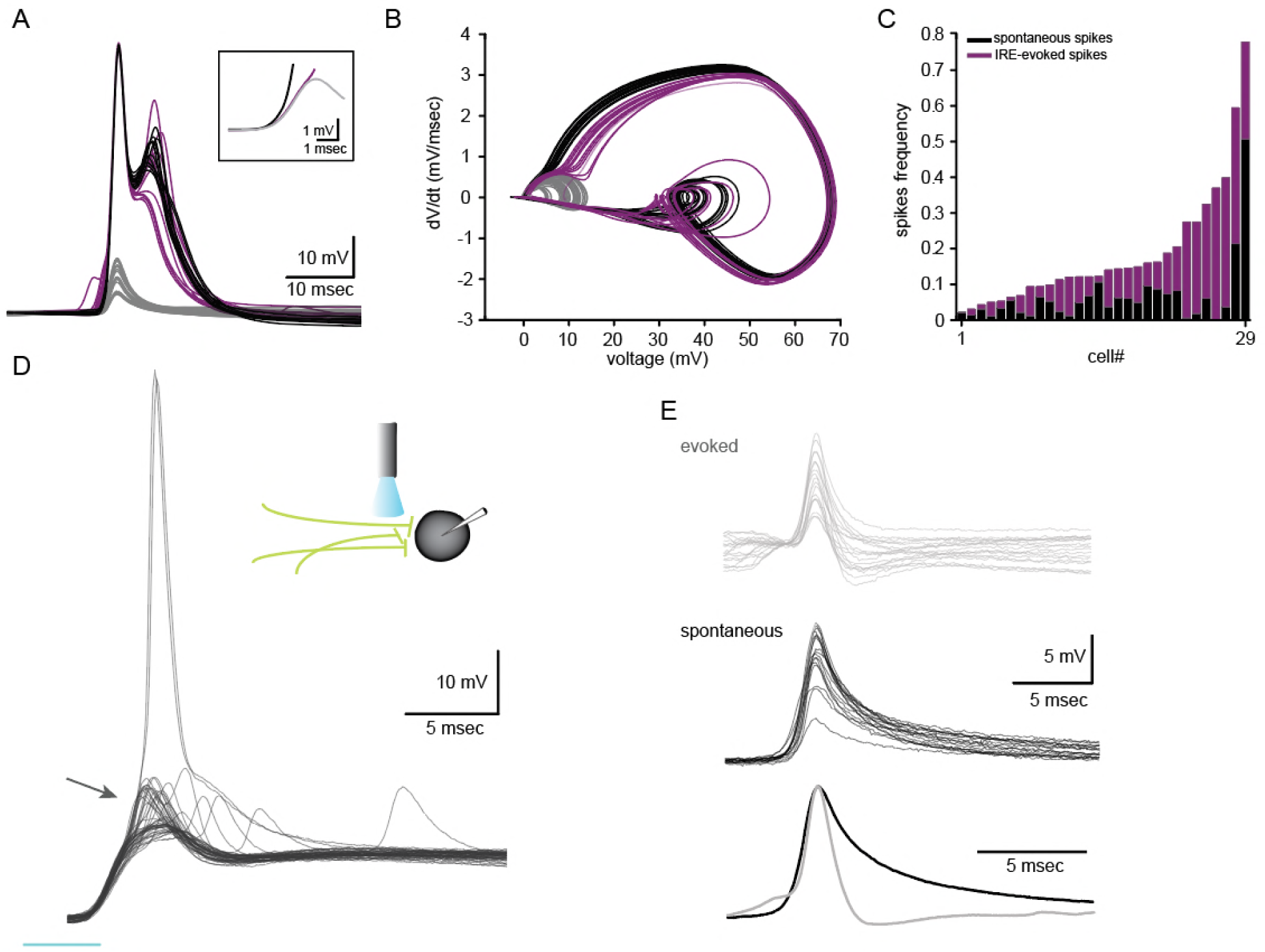
Internal Regenerative Events (IREs) are the major source of action potentials and can be evoked by synaptic inputs. (A-B) Voltage (A) and phase plots (B) of spontaneous occurring IREs and action potentials. Purple - action potentials that were triggered by IREs, gray – IREs that failed to trigger an action potential. Black - fast rising action potential. (C) frequency of spontaneous action potentials (black bars) and of IRE-evoked action potentials (purple bars) in 29 neurons. (D) Thy1-ChR-EYFP mice were used to optogenetically activate incoming excitatory axons (inset). Superimposed traces recorded from an olivary neuron in response to 5 ms light pulse, showing synaptic potentials topped with IREs (arrow) that occasionally triggered action potentials. (E) Upper panel: IREs evoked by the synaptic input. Middle panel: spontaneous IREs recorded from the same neuron. Lower panel: normalized average of the spontaneous (black) and the evoked (gray) events. The evoked IREs were isolated by subtracting the average subthreshold synaptic waveform.

We then examined the possibility of activating these events by synaptic inputs, using Thy1-ChR2-YFP transgenic mice in which ChR2 is expressed in axons that deliver excitatory inputs to the IO but not in IO neurons (Lefler, Torben-Nielsen and Yarom, 2013). As shown in Figure 4D, a short light pulse (inset) triggered a compound response comprised of synaptic potentials crowned with fast events that occasionally triggered action potentials. The evoked events were isolated by subtracting the synaptic response (Figure 4E, top panel, gray traces). They were then compared to IREs that occurred spontaneously (middle panel, black traces) by averaging and normalizing each of the groups (lower panel). Whereas the rise time for the spontaneous IREs matched that of the evoked IREs perfectly, the decay of the evoked response was considerably faster. This difference might be the result of a depolarized membrane potential during the synaptic event, which is in line with the effect of DC depolarization on the IRE waveforms (Figure 3), as well as the result of high membrane conductance.

These results strongly support the possibility that the fast events are generated within the neuron either spontaneously or in response to synaptic input. Their fast kinetic suggests that sodium current is involved; in fact, these fast events were not detected in 27 neurons that were injected with QX-314 (2-5 mM). This contrasts with normal conditions in which only 28% of the cells were devoid of fast events. Furthermore, in a complex experiment with two patch electrodes in the same neuron, one filled with regular intracellular solution and the other with QX (Supplemental information, Figure S1), we found that in the control condition both IREs and spikelets were readily detected (Figure S1A) whereas when breaking into the cell with the QX-containing electrode, the fast events could not be detected whereas the spikelets were almost unchanged (Figure S1B). Finally, in CX 36 knockout mice, where gap junctions were eliminated (De Zeeuw *et al.*, 2003), fast and high amplitude events could still be encountered (Figure S1C). These results strongly confirm our supposition that the fast events are generated within the neuron.

### Modelling the Internal Regenerative Events (IREs)

Our experimental observations suggest that IREs represent regenerative responses within the recorded neurons. It was suggested that such responses could result from either dendritic ‘hot spots’ (Spencer and Kandel, 1961) or from the failure of antidromic axonal spikes at remote Nodes of Ranvier (Traub, Colling and Jefferys, 1995; Avoli, Methot and Kawasaki, 1998). The most characteristic feature of the IREs is that for any given cell, their amplitude clearly segregates into distinct groups, whereas their waveform is essentially identical (Figure 1C-D). We used a modeling approach to explore the conditions in which this feature could be reproduced, either for dendritic ‘hot spot’ or for axonal spike failure.

To examine the possibility of “dendritic hotspots” we developed detailed compartmental models of three 3D reconstructed olivary neurons; a sample modeled cell is shown in Figure 5A-C. To simulate “hot spots”, we injected, at every dendritic location, a simulated spike (Stuart and Sakmann, 1994) with a 100 mV amplitude and a 2 ms half-width (Figure 5D). We searched for the dendritic locations where the resultant somatic voltage response resembled the experimental IREs, with half-widths ranging from 3.25 to 3.5 ms (see Figure 1F). This yielded nine possible confined dendritic locations in the modeled cell (colored dendritic segments in Figure 5C). The respective simulated responses are shown in Figure 5E. Thus, although the amplitudes of these responses varied from 2 mV to 6.5 mV (Figure 5E), their waveforms were almost identical (Figure 5F). Similar results were found in two other modeled cells, thus demonstrating that only well-placed dendritic “hot spots” could replicate the experimental results.

**Figure 5.**
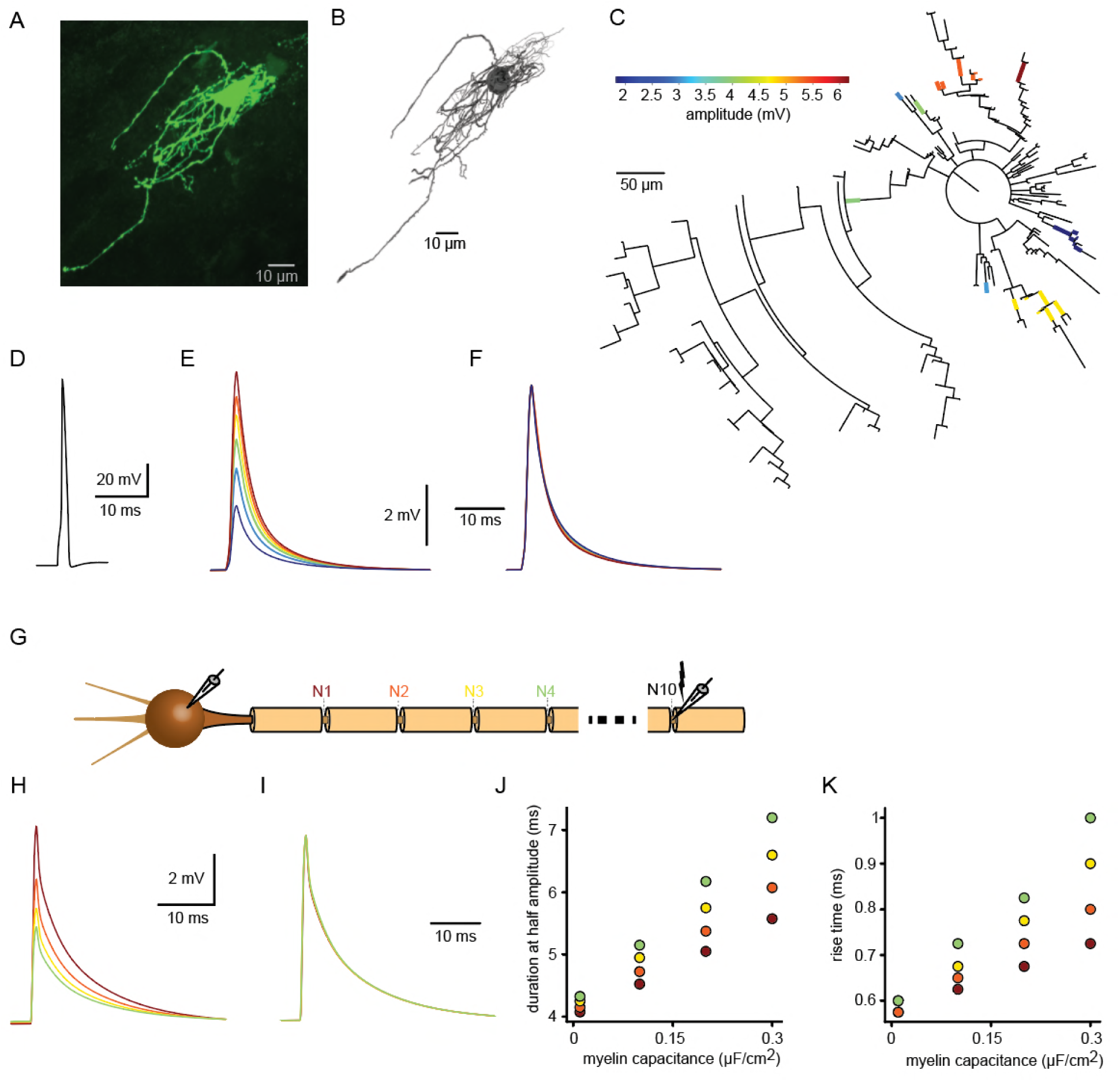
Dendritic hotspot (A-F) and axonal failure (G-K) models for the generation of internal regenerative events. (A) Confocal image of a Biocytin filled olivary neuron. (B) 3D reconstruction of the neuron in a. (C) Dendrogram of the reconstructed neuron with possible ‘hot spot’ locations color coded according to the amplitude of their somatic response (see color bar). (D) the action potential waveform generated in each of the hot spots. (E) The somatic responses to the activation of each hot spot. (F) Normalization of the responses in E. (G) Schematic Illustration of the model consisting of a soma, an axon hillock, an axon initial segment and 10 myelinated segments, separated by active nodes (N1-N10). (H) The somatic responses to failure of the antidromic action potential in N1-N4 (color-coded). (I) Normalization of the responses in H. (J-K) Duration and rise times of the events plotted against the myelin capacitance.

To examine the possibility of axonal spike failure we constructed a model of a myelinated axon connected via an initial segment to an isopotential soma connected to several passive dendrites (Figure 5G; see Methods). We examined the somatic voltage response to an antidromic action potential (activated by short current injection at node 10) during successive blockade of the Nodes of Ranvier (starting with node 1 only, then node 1+2, etc.). As more nodes are blocked, a smaller somatic response is expected (Figure 5H, brown, red, yellow and green traces, respectively). Surprisingly, normalizing the amplitude of these responses (Figure 5I) revealed that the waveform was almost completely unaffected by the location of the propagation block. This was due to the small effective capacitance of the myelin. Indeed, increasing the myelin capacitance enhanced the difference between the waveforms of the somatic responses (Figure 5J-K).

Both models reproduced the characteristic features of the IREs; i.e., identical waveforms and variable amplitudes. However, it is difficult to envisage a biological mechanism that either specifically localizes channels in a restricted dendritic “hot spot” or that simultaneously blocks two, three or more Nodes of Ranvier. Interestingly, however, olivary neurons have unique axons which provide a third possible explanation for the IREs. The classical work by De Zeeuw and colleagues (De Zeeuw *et al.*, 1990) demonstrated the existence of a spine apparatus in the axonal hillock of olivary neurons. We modeled these axonal spines while taking their different neck lengths into account (Figure 6A) and assumed that their head membrane is excitable (see Methods). Figure 6B shows the somatic responses, ranging from 6 to 9 mV, that resulted from action potentials generated in these axonal spines by a short current injection. The somatic responses had identical waveforms (Figure 6C). Importantly, these axonal spines are innervated by excitatory synapses (see De Zeeuw *et al.*, 1990, Figure 9C), enabling their individual activation. We next examined whether the model of excitable axonal spines could replicate the experimental finding that the IREs exhibit an increased duration upon membrane hyperpolarization (Figure 3). The results confirmed that injection of hyperpolarizing or depolarizing currents into the model soma altered the action potential waveform in the spine head (Figure 3D), particularly the after-hyperpolarizing phase, which, as expected, turned into a depolarizing phase (black arrow in Figure 6E, left panel). As a result, the duration of the somatic response increased (red arrow in Figure 6E, right panel). This effect was observed for three levels of somatic current injections (−150 pA, -30 pA and +30 pA) for all four spines (Figure 6F). The normalized responses are displayed in Figure 6G, and illustrate the prolongation of the repolarizing phase of the somatic responses with hyperpolarization, as found experimentally.

**Figure 6.**
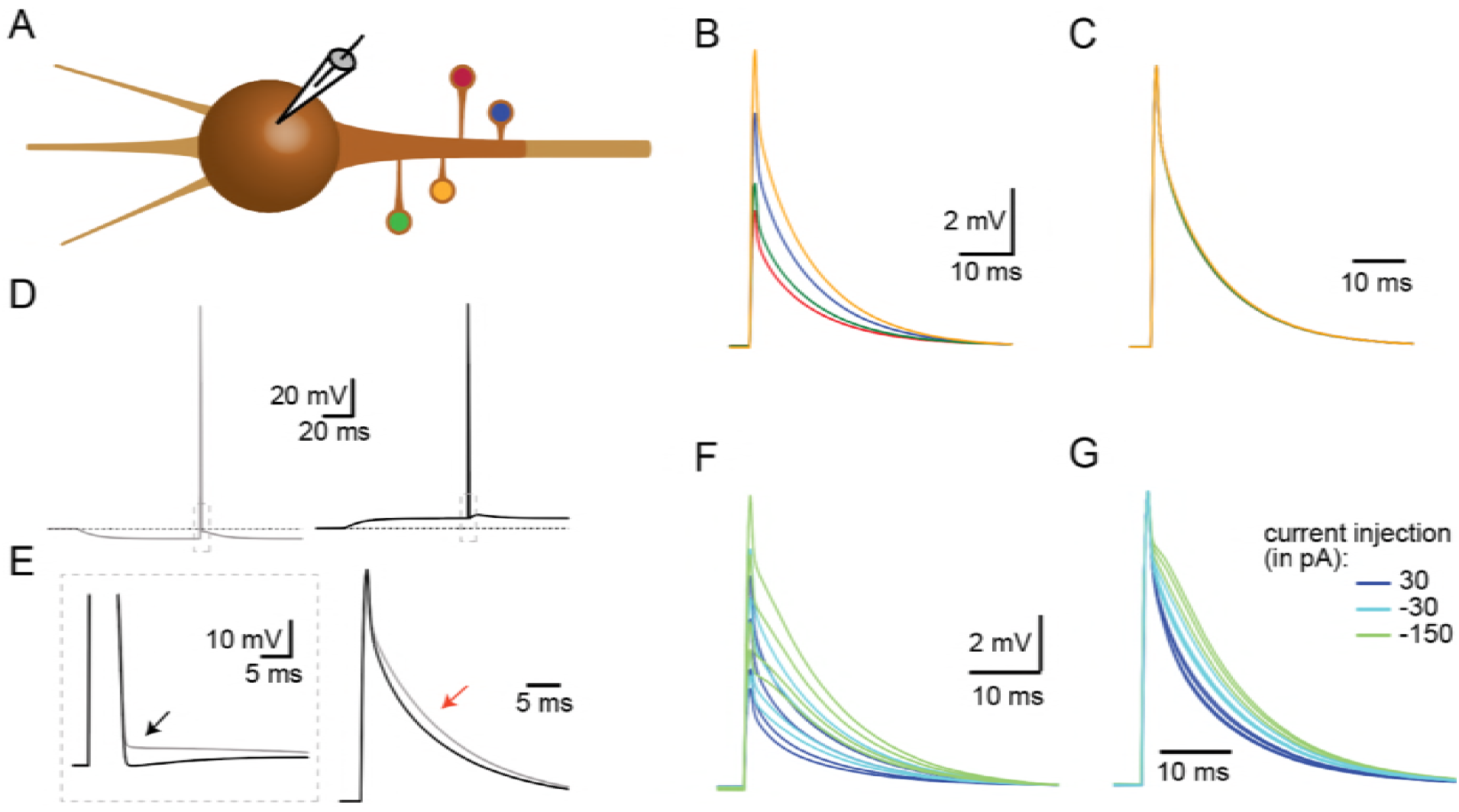
Initial segment spine model for the generation of internal regenerative events. (A) Schematic representation of the model where 4 different spines are located along the axon hillock. (B) Somatic response to action potentials generated in the spine head of the spines (color coded in A). (C) Normalization of the responses in B. (D) The effect of somatic hyperpolarization (gray trace, left panel) and depolarization (black trace, right panel) on the action potential waveforms. (E) left panel, Superposition of the dashed rectangles of the action potentials in d and e showing the difference in the after-hyperpolarization (black arrow) between hyperpolarization and depolarization. Right panel, Superposition of the somatic responses showing the prolongation of the response with hyperpolarization (red arrow). (F) Somatic response evoked by spine action potentials at three levels of somatic DC current injection. (G) Normalization of the responses in F.

### Estimating network architecture from dual cell recordings of simultaneously occurring spikelets

Figure 7A depicts the spontaneous activity recorded simultaneously from two neurons. As described above (Figure 2) action potentials (diagonal bars) that occurred irregularly in either of the two neurons were always associated with spikelets in the paired neuron (Figure 7B). The subthreshold activity, which was dominated by spikelets as well as IREs, appeared randomly in the two neurons. However, occasionally spikelets occurred simultaneously in both cells (marked in Figure 7A and shown at high resolution in Figure 7C) which we refer to as ‘common spikelets’.

**Figure 7.**
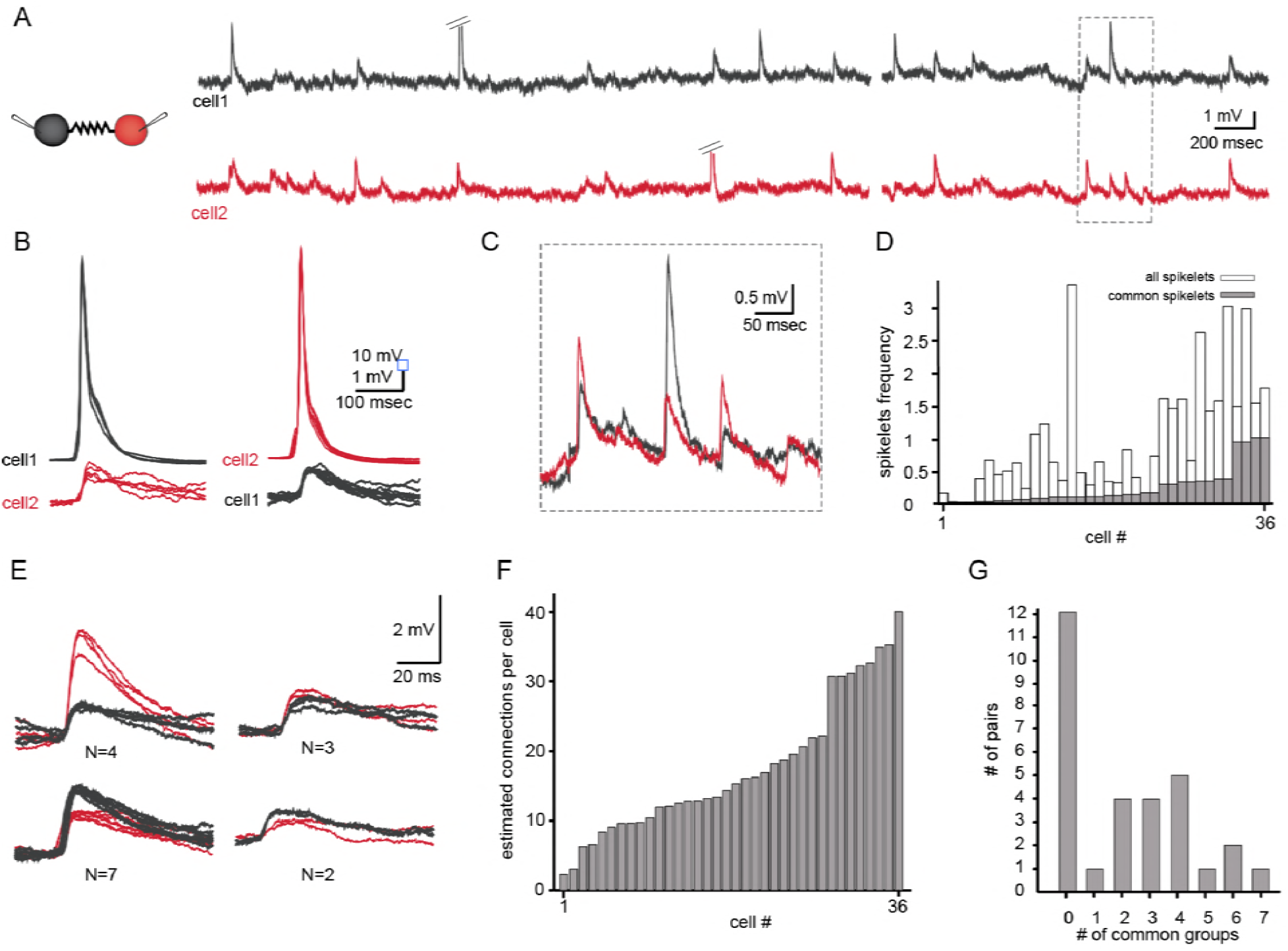
Common groups of spikelets during paired recording reveals the average number of neurons that are electrically coupled to each neuron in the slice. (A) Simultaneous recording from two electrically coupled neurons. Action potentials were truncated (doubled diagonal lines) and an example of the occurrence of common spikelets is marked (dashed rectangle). (B) Superimposed traces of spontaneous action potentials in either cell 1 (black neuron) or cell 2 (red neuron) and the corresponding spikelets in the other neuron. (C) Higher magnification of the rectangle marked in A, showing spikelets that occur simultaneously in both neurons. (D) Histogram of the frequency of spikelets in neurons recorded in pairs, showing all the spikelets (white bars) and all the common spikelets (gray bars; n=18 pairs). (E) Example of common spikelets from the pair presented in a-c. The spikelets could be divided into 4 groups, with N = 2-7 spikelets in each group. (F) Histogram of the estimated number of neurons that are electrically coupled to each of the pair-recorded neurons (n=18 pairs). (G) The distribution of the number of common groups in all recorded pairs.

Each of the three examples shown in Figure 7C, which occurred without measurable time difference, have variable amplitudes. The first and the third spikelets had larger amplitudes in the red neuron (cell 2) whereas the middle spikelet had a larger amplitude in the black neuron (cell 1). Since action potentials in one neuron evoke very similar spikelets in the other (Figure 2A-B), the most likely explanation is that each of these common spikelets represents the action potential in an additional neuron that is coupled to both of the recorded neurons (see Discussion). On the population level, 18 out of 38 pairs had common spikelets (47%). Of these pairs, the occurrence of common spikelets varied from 0.02 to 1.1 /sec, which is 3.5-66% of the total number of measurable spikelets (Figure 7D).

The occurrence of common spikelets can be used to estimate the number of neurons that are electrically coupled to each neuron in the olivary network. In this example of a paired recording, 4 different groups were identified by clustering the detected common spikelets by the amplitude value and amplitude ratio between the two neurons (Figure 7E), which indicates that at least 4 neurons were electrically coupled to both recorded neurons. Further analysis of these data provided an estimate of the total number of neurons connected to each of the 2 recorded neurons: in this example, 16 common spikelets in 4 groups were recorded; in other words, these neurons fired on average 4 times during the recording period. Since 25 and 50 spikelets were recorded overall for the black and red neuron, respectively, the 9 and 32 non-common spikelets argue for the presence of additional electrically coupled neurons. Assuming that all the neurons fired at similar rates (∼4 spikes during the recording period), these non-common spikelets thus represent spikes in 2 additional neurons connected to the black neuron and 8 neurons connected to the red neuron. The result of this numerical consideration is that the black neuron is connected to the red neuron, to 4 additional neurons that are connected to both recorded neurons and an estimated 2 additional neurons, thus totaling 7 neurons. Similarly, the red neuron is connected to up to 13 neurons. This analysis was performed on 18 dual recordings and the results, which are summarized in Figure 7F, indicate that a neuron can connect to as many as 40 other neurons (average of 19.2±10.3). It should be noted that the use of a slice preparation undoubtedly contributed to the wide range of connected neurons and to some degree of underestimation (see Discussion).

Further insights into the organization of the network can be extracted from the distribution of the number of groups of common spikelets. As shown in Figure 7G, the number of common groups varied from 0 to 7 with a likely higher incidence at 2 - 4 groups. As will be shown below, this type of distribution cannot be the result of random connectivity where the probability of connection depends solely on the distance between the recorded cells.

We re-examined the approach we used to estimate the number of connections per neuron by reconstructing a realistic olivary network (Figure 8A, see Methods). The firing rate of neurons in the network was set to 0.058 Hz ± 0.04 Hz (as observed experimentally) and the number of common spikelets in pairs of neurons occurring within 15 min of simulation was measured. Recordings from a sample pair are shown Figure 8B. In this example, four groups of spikelets that appear 26, 16, 65 and 32 times were detected (Figure 8B). By applying the same calculation as performed in the experimental observations (Figure 7), we concluded that the red neuron was electrically connected to 24 neurons whereas the black neuron was connected 26 neurons. In this model, the red and black neurons were actually connected to 17 and 20 neurons, respectively (pink circles in Figure 8C). We performed the same calculation in 20 randomly selected pairs of neurons and plotted the estimated versus the real number of connections per cell (Figure 8C). The results were distributed along the diagonal, (with the average marked by + sign), thus demonstrating the validity of this approach in estimating the number of connections for each neuron. However, the accuracy of the estimation depends strongly on the variability in firing rate and the recording duration; the accuracy decreases the shorter the recording duration and the higher the firing rate variability, but the error in estimating the average number of connections for each cell was still small (Supplemental information, Figure S2, see discussion).

**Figure 8.**
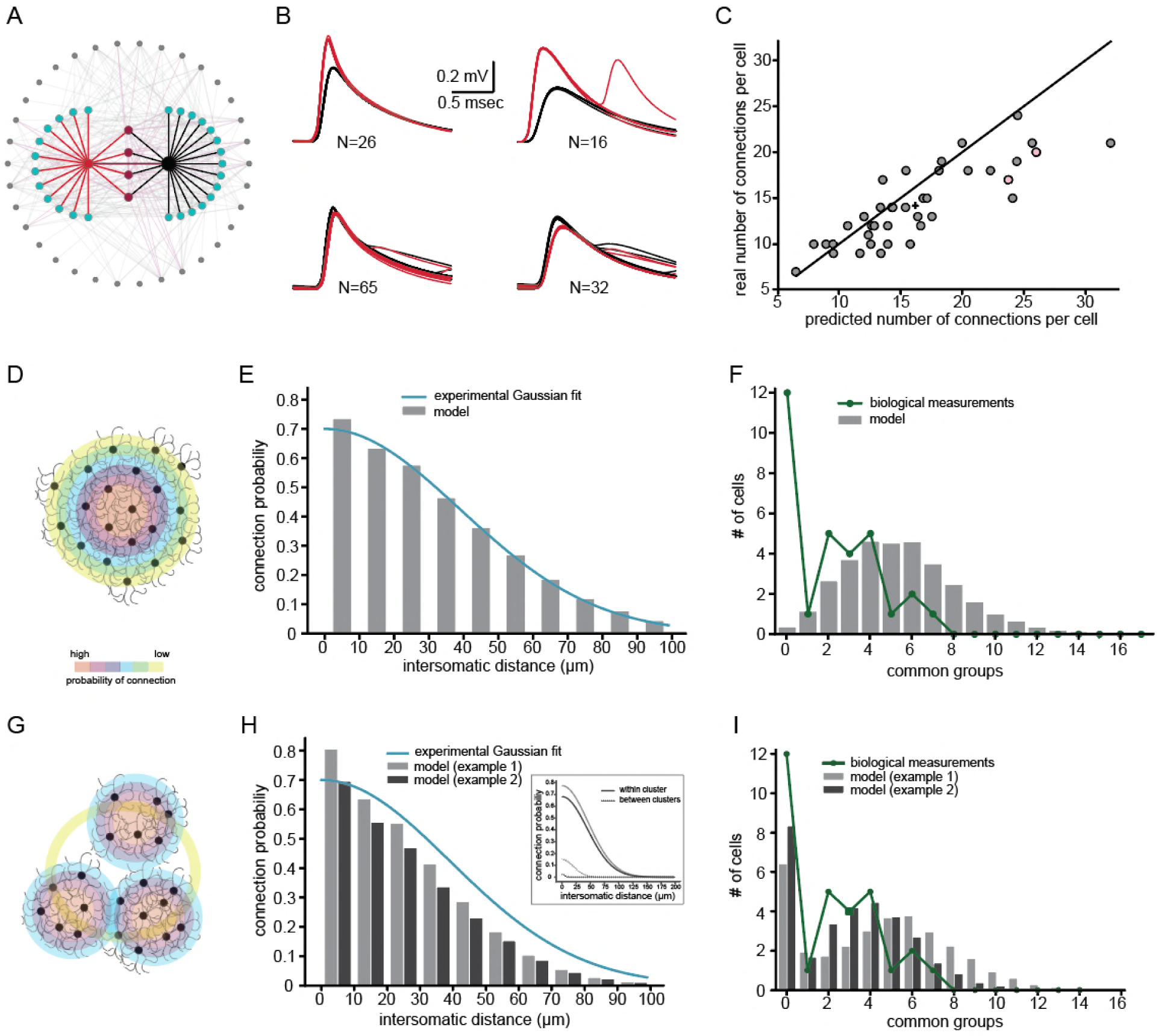
Simulations examining both the method used for estimating the number of connections per cell and the network connectivity that accounts for the experimental distribution of common groups of spikelets. (A) Schematics of the modeled network where the recorded pairs of neurons (black and red circles) are connected to four common neurons (purple) and to 12 and 15 additional neurons (cyan); 34 other neurons that are connected to either the cyan or the purple neurons are also shown (gray). (B) the four common groups of spikelets recorded in the black and red cells in A. (C) Plot of the predicted number of connections per cell, estimated from the common groups of spikelets, against the real number of connections per cell. The line marks the diagonal, the + sign marks the mean and the pink circles represent the two cells in A. (D-F) the expected distribution of the common groups in a model where the probability of connection is distance-dependent. (D) Schematic illustration of a distance dependent connectivity. The connection probability is color coded. (E) The probability of connection in the model (gray bars) and in the experiments (blue line) as a function of the inter-somatic distance. The blue curve represents a Gaussian fit to the data. (F) Distribution of the common groups in the model (gray bars) and experiments (green line) for cells of up to 40 um apart. (G-I) Same as D-F for two networks that are organized in clusters of neurons with a high probability of connection within a cluster (continuous line in inset; matching color-code as the bars in H and I) and a low probability between clusters (dashed lines in inset) in both networks. Each cluster consisted of about 40 neurons.

The experimental distribution of the number of groups of common spikelets (Figure 7G) provides additional information on the organization of the network. We examined two possible connectivity patterns that might support this type of distribution (see Methods; network connectivity matrices). The first is a network where the probability of a connection between two neurons depend solely on their inter-somatic distance (Figure 8D). The second assumes that the network is organized into clusters of neurons (Figure 8G), where the connection probability within a cluster is larger than between clusters (both probabilities are distance-dependent). We searched for the parameters that yielded the best fit to the experimental observations; namely, the distance-dependent connection probability (see Devor and Yarom, 2002a and Figure 8E, H) and the distribution of common neighbor (proxy for groups of common spikelets in Figure 8F, I). Our search yielded an interesting result. Whereas the simple distance-dependent network captured the distance-dependent probability of a connection (Figure 8E), it failed to reproduce the distribution of common groups as found experimentally (Figure 8F). On the other hand, when the modeled networks were organized in clusters, it replicated both the experimental distribution of common groups (Figure 8I) as well as the distance-dependent connection probability (Figure 8H). Note that in all modeled networks, each neuron was connected to about 11-21 neurons, which lay within the numbers estimated from the experimental observations (Figure 7F).

Figure 8 also shows that the experimental results could be reproduced in networks with various distance-dependent connectivity profiles (see Methods), provided that the network is organized into clusters of neurons. Figure 8 H-I presents two examples of networks that differ in their within- and between-connectivity profiles (black vs. gray lines in Figure 8H inset). The resultant distance-dependent connectivity and the distribution of the common groups for these two cases are denoted by the black and gray bars. Thus, it seems likely that the inferior olive is organized into clusters of neurons with a higher probability of connection within clusters and a low connection probability between clusters.

## Discussion

In this study, we measured subthreshold unitary activity from neurons of the inferior olive in slice preparation. The implementation of a variety of experimental approaches linked with computational simulation led to several important conclusions regarding the organization of the network of electrically coupled neurons in the inferior olive nucleus. We showed that there are two populations of non-synaptic unitary events that differ in their waveform and amplitude. The spikelets represent the occurrence of action potentials in a coupled neuron, and the IREs are likely to represent intrinsic regenerative responses at remote locations. It should be noted that in the scientific literature the word ‘spikelets’ refers to all non-synaptic subthreshold unitary events and thus creates a certain lack of consistency regarding their origin (see Introduction). The uniqueness of our experimental system lies in its ability to differentiate between two types of non-synaptic unitary events and thus to characterize them. We then used the spikelets recorded simultaneously from two neurons to gain insights into the size and organization of the electrically coupled network within the IO nucleus. Analysis of the experimental results showed that each olivary neuron is connected to ∼ 20 other neurons and theoretical considerations indicate that the nucleus is organized into clusters of neurons, where the probability of connection within a cluster is higher than the probability of connection between clusters.

### The Internally Regenerative Events

These fast subthreshold responses are a mystery. At first glance, they appeared to be a representation of an action potential in a coupled cell, and the difference from spikelets could be accounted for satisfactorily by direct versus indirect coupling where the coupling is mediated through an intermediate neuron. The simulations showed that this can explain the different waveforms as well as the amplitude of the two types of responses (see Supplemental information, Figure S3). However, out of over 45 paired recordings we **never** encountered a pair where an action potential in one neuron elicited an IRE in the coupled neuron. Furthermore, an IRE in one neuron never coincided with a spikelet in the other neuron. Moreover, whereas the probability of spikelets was independent of the membrane voltage, the IRE showed an increased probability at depolarization (Figure 3). Finally, IREs were recorded from the olivary neuron in CX 36 knockout mice (De Zeeuw *et al.*, 2003)(see Supplemental information, Figure S1). Thus, we concluded that the large unitary events represent localized regenerative responses at remote cellular locations.

The locations and mechanism of generation of the IREs are not completely resolved. A localized regenerative response requires a localized concentration of voltage-dependent channels that can support the generation of a local action potential. We investigated three possible locations: ‘hotspots’ along the dendritic tree, spike generation at different Nodes of Ranvier along the axon, and spike generation at spines located along the axon initial segment. All three possibilities can support the occurrence of unitary events of different amplitudes and almost identical waveforms and can account for the voltage dependency of the waveform. However, the locations of the dendritic hot spots, which can support similar waveform with different amplitudes, are restricted to rather small dendritic segments distributed on different dendritic branches without any common structural features. Thus, it is difficult to envisage a sophisticated mechanism that determines the location of the voltage-dependent ionic channels within a rather large dendritic tree that enables the generation of action potentials **solely** at the designated locations. Finally, a full action potential at the dendritic tree should be able to propagate through the gap-junctions to the post junctional neurons and thus should be revealed during paired recordings.

The Nodes of Ranvier, on the other hand, are well-organized sites where the locations of the voltage-dependent ionic channels are restricted to small patches organized along the axon at similar intervals (Murtie, Macklin and Corfas, 2007). This type of organization is well suited to generate, at somatic level, a set of subthreshold events organized in well-defined groups of different amplitudes. The almost identical waveform of such events reflects the very low capacitance of the axon (due to the myelin sheet), which prevents waveform changes. However, to account for the failure of axonal propagation, one has to assume that action potentials can be generated down the axon away from the cell body (Pinault, 1995; Sasaki, 2013). Moreover, the fact that the axon, or more likely the axon initial segment, is usually regarded as the site with the lowest threshold (Palmer and Stuart, 2006), cannot be readily applied in our case, since simultaneous failure at multiple nodes are needed to generate different amplitudes.

Finally, we were able to simulate the experimental observation assuming that the source of the IREs are the spines that emerge from the axon initial segment. If these spines can support action potentials, as suggested in other systems (Segev and Rall, 1998; Araya *et al.*, 2007; Bywalez *et al.*, 2015), it should appear at the somatic level as an IRE. In our model, the different amplitudes mostly reflect the neck length. In fact, the detailed ultrastructural study by De Zeeuw and colleagues (De Zeeuw *et al.*, 1990) indicated that up to 8 spines emerge from the axon initial segment of some olive neurons. Furthermore, these spines are innervated by chemical synapses, where the specific activation of individual spines are bound to occur. Thus, it is more likely that the source of the IREs are these axonal spines.

Spines emerging from an axon is not a common feature of central neurons. As we demonstrated, IREs are strong enough to activate the neurons (Figure 4) and thus generate a unique situation where the activation of the neuron is independent of the background synaptic activity that takes place in the soma and the dendritic tree. It is tempting to speculate that this arrangement ensures activation of the neurons in a state of “emergencies”. Thus, when the system is in need of a fast response, it activates these inputs that bypass the ongoing activity, thus insuring an immediate response.

### The electrically coupled network in the olivary nucleus

*Spikelets and subthreshold oscillations:* The formation of electrically coupled network within the olivary nucleus is well-established. Here we suggested that this network, operating as a synchronous rhythmic device, is capable of generating precise temporal patterns (Jacobson, Rokni and Yarom, 2008). Both synchronicity and rhythmicity are generated by the delicate interplay between electrical coupling and ionic conductances. Thus, a single cell by itself cannot oscillate, whereas in a network formation the neurons generate subthreshold oscillations (Manor *et al.*, 1997; Loewenstein, Yarom and Sompolinsky, 2001; Chorev, Yarom and Lampl, 2007). In this work, we studied the relationship between spikelets and subthreshold oscillatory activity and found that in oscillating cells the occurrence of spikelets coincided with the depolarizing phase of the oscillation whereas in non-oscillating cells they seemed to be randomly distributed (Figure 2). This result strongly supports our previous hypothesis that the occurrence of subthreshold oscillations is a network phenomenon. Therefore, when the recording cell is oscillating, the entire network is synchronously oscillating, thus generating action potentials at the peak of the oscillation that appear in the recorded cell as spikelets. Theoretically, by calculating the number of spikelets at the peak of the oscillatory activity, one should be able to calculate the number of coupled neurons in the oscillating network. Although tempting, this is practically impossible because spikelets, given their small amplitude and noisy environment, cannot be classified into groups. Therefore, we used the common spikelets to estimate the number of coupled neurons. *Common spikelets:* Recording from two olivary neurons revealed spikelets that occurred simultaneously in both recorded neurons. Given that the average rate of spikelets is 0.56±0.62Hz, the probability that these common spikelets reflect random occurrence is extremely low. Furthermore, the repeated appearance of common spikelets with the same relationships (amplitude ratio and waveforms) further supports the non-random occurrence of these events. Thus, the accurate timing of common spikelets can only be attributed to a common source; i.e., a single pre-junctional neuron. The number of neurons that were coupled to the two recorded neurons, which varied from 1 to 7, should be correlated with the size of the network; more common spikelets are expected in larger interconnected networks.

Our simple method of calculating the size of a coupled network is based on data obtained during simultaneous recordings from two neurons and on the assumption that the neurons display a similar rate of spontaneous spiking activity. Our simulations show that the accuracy of this method is mainly affected by variability in the firing rate (Supplemental information, Figure S2). To minimize the error in estimating the firing rate of the neurons in the network, we used only the spontaneous rate of the common spikelets, and not the firing rates in the recorded neurons that are affected by the intracellular recordings. Under this assumption, we showed that neurons are connected to 3-40 other neurons. This is in line with other studies reporting 1-38 (Hoge *et al.*, 2011) or 0-33 (Placantonakis *et al.*, 2006) dye coupled neurons. This broad variability can be attributed to the use of an in-vitro system, where differences in the number of cells and the integrity of the circuit are characteristic features. Alternatively, this large variability might reflect an innate feature of the IO nucleus where electrical coupling is under continuous modulation (Lefler, Yarom and Uusisaari, 2014; Mathy, Clark and Häusser, 2014; Turecek *et al.*, 2014).

In addition to calculating the number of connected cells, the distribution of common spikelets enabled us to study the connectivity profile within the nucleus. Our data showed that each two neurons had 1-7 common groups, and a normal distribution with an average of 3-4 groups. However, there was a large number of paired recordings that failed to show common groups. Using a theoretical approach, we demonstrated that this distribution should not be expected if we assume that the probability of connection depends solely on the distance between the neurons. On the other hand, if the nucleus is organized into clusters where the probability of connection within the cluster is higher than between clusters, the observed distribution of common groups can be reproduced. Although the size of the clusters, as well as the probability of connection cannot be defined with the current data, this constitutes the first physiological study that supports the assumption of clustered organization of the nucleus deduced mainly from morphological studies.

In summary, we presented a comprehensive study that implemented a wide range of research approaches to unravel the functional architecture of the inferior olivary network. We showed that by analyzing spontaneous subthreshold events, new insights into the organization of the network can be gained, thus paving the way for a novel experimental and theoretical approach to the study of electrically-coupled networks.

## Acknowledgments

This study was supported by the Israel Science Foundation (http://www.isf.org.il/) grant #1496_2016. I.S and O.A were supported by grant agreement no. 604102 ‘Human Brain Project’, a collaborative grant under the Blue Brain Project and a grant from the Gatsby Charitable Foundation. Y.L. was supported by a postdoctoral fellowship from the Edmund and Lily Safra Center for Brain Sciences (ELSC).

## Author contributions

These authors contributed equally: Y. Lefler and O. Amsalem

Experimental work and data analysis were conducted by Y.L, and computational work and data analysis by O.A. All authors interpreted the data and wrote the manuscript.

## Materials and Methods

### Animals

All experimental procedures were approved by the Hebrew University’s Animal Care and Use Committee. Brain stem slices were prepared from the following strains of mice: C57BL/6, B6.Cg-Tg (Thy1-COP4/EYFP)(Jackson Laboratory) and Gad2-tm2(cre)Zjh/J (Jackson Laboratory).

### Slice preparation

Mice were anesthetized with an intraperitoneal injection of Pentobarbital (60mg/Kg), and 300 µm coronal brainstem slices containing the inferior olive were dissected using a Campden 700smz slicer (Campden Instruments), in 35°c physiological solution containing 126 mM NaCl, 3 mM KCl, 1.3mM MgSO_4_, 1.2mM KH_2_PO_4_, 26mM NaHCO_3_, 10mM glucose, and 2.4mM CaCl_2_, gassed with 95% O_2_ and 5% CO_2_. Slices were left in physiological solution at 35°c for 0.5 - 8 hours until transferred to the recording chamber.

### Electrophysiological recordings and ChR stimulation

The recording chamber was perfused with 95% O_2_ and 5% CO_2_ physiological solution at 24°c– 28°c. Slices were visualized using a 40X water-immersion objective in an Olympus BX61WIF microscope equipped with infrared differential interference contrast (DIC). In order to record from intact olivary networks, recordings were targeted to the deepest neurons possible in the slice. Whole-cell recordings were performed using 6-9 MΩ glass pipettes with intracellular solution containing 4mM NaCl, 10^-3^mM CaCl_2_, 140mM K-gluconate, 10^-2^ mM EGTA, 4 mM Mg-ATP, and 10 mM HEPES (pH 7.2). Signals were acquired at 10-20KHz using a Multiclamp 700B (Molecular Devices) and LabView-based custom-made acquisition software (National Instruments and ZerLabs). For the ChR experiments, a custom -made digital mirror light stimulator with a LED light source (460 nm; Prizmatics) was used to activate the ChR at defined locations on the slice. In some experiments either TEA (10 mM), CNQX (10-40 µM), DNQX (20-40 µM) and/or AP-5 (40-100 µM) were added to the recording solution or QX-314 (2-10 mM) was added to the pipette intracellular solution.

### Reconstructing neurons

In a few experiments (using C57BL/6 mice), Neurobiotin (0.5%; Vector Laboratories) was added to the pipette solution to label the recorded neurons. Slices were then fixed in 4% paraformaldehyde overnight, washed in PBS and stained with 1 µg/ml Streptavidin AlexaFluor 488 (Life Technologies).

### Data analysis and Statistics

Analysis was performed using MATLAB (R2014b and R2016a, MathWorks) for the experimental data and Python 2.7 for the simulation data. The 71 neurons that were selected for detailed analysis had a frequency of subthreshold events exceeding 0.02 Hz. The events were divided into two different groups according to their amplitude and rise time. The event rise time was calculated as 10%–90% of the amplitude. The IRE groups were clustered using the K-means clustering method. The coupling coefficient (CC, Figure 2) was calculated as the ratio between the change in the steady-state voltage of the post-junctional cell and that of the pre-junctional cell in response to 250 ms current injection in the pre-junctional cell. The spike coupling coefficient was measured as the amplitude of the post-junctional spikelet divided by the amplitude of the pre-junctional action potential. A Pearson correlation was used to calculate the p-value of the linear regression in Figure 2C. The Inter-spikelet-interval (ISLI) in oscillating neurons (Figure 2G) was only calculated in neurons that had more than 150 spikelets during the session. The ISLI histogram was computed using 4 ms time bins, and the autocorrelation in oscillating neurons was calculated using a lag of 1 ms.

The effect of the DC current injection in (Figure 3 C-H) was measured in 16 neurons for different values of current injection. For each neuron, the average value for each current injection was calculated (gray dots in Figure 3C-H) and fitted with a linear line (dashed gray lines). To calculate the average slope (black lines), we averaged the gray dots for each current injection (black dots) and fitted them with a linear line. Error bars represent STD. A one-sample t-test was used to compare the distribution of the slopes of the linear fits of each cell (dashed gray lines) to a distribution with a mean equal to zero. A paired t-test was used to test for differences in the effect on spikelets and IREs. The frequency values for spikelets and IREs (Figure 3G-H) were normalized for each cell to the highest value. The normalized IREs frequency (Figure 3H) was calculated from both IREs and the action potentials that were evoked from IREs.

To define spikes that were evoked from IREs (Figure 4A-C), we examined the phase plot of all spikes in each cell and manually searched for a clear gap between the curves around the threshold for spike initiation. To isolate the ChR-evoked subthreshold events (Figure 4E), synaptic responses that did not evoke IREs or spikes were averaged and subtracted from individual responses with IREs.

Common spikelets (Figure 7) were defined as spikelets that were detected in a paired recording in both cells simultaneously. Groups of common spikelets were clustered using k-means analysis on the peak voltage, followed by manual curation in which we verified that all common spikelets in a group had a similar shape. To estimate the number of connections per cell, the total number of spikelets (T_spikelets_) was multiplied by the number of groups (N_groups_) and divided by the number of common spikelets (C_spikelets_):

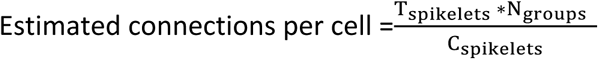

If the two recorded cells were coupled (i.e., a spike in one cell gave rise to a spikelet in the other cell), +1 was added to the estimation of connections for these two cells.

### Neuron models

Using the Neurolucida software (MBF Bioscience), three olivary neurons were reconstructed from fluorescence image stacks acquired using a Leica TCS SP5 confocal microscope (Leica Microsystems). To compensate for tissue shrinkage, the z-axis of the reconstruction was multiplied by a factor of 3. A compartmental model was generated from the morphological reconstruction using NEURON (Carnevale and Hines, 2006). The axial resistance (R_a_) was set to 100 Ωcm, the specific membrane capacitance (C_m_) to 1 µF/cm^2^ and the specific membrane resistivity (R_m_) for the three reconstructed cells were 4300, 4500, 3800 Ωcm^2^ respectively. These values were chosen to yield an input resistance (R_in_) that was within the experimental range (115±43 MΩ).

### Axon model

The model was based on a previous axonal model (Chen *et al.*, 2002) (ModelDB, accession number 3793) with some modification. The model had the following compartments: soma, five cylindrical dendrites, an axon hillock (AH), an axon initial segment (AIS), ten myelinated segments and ten Nodes of Ranvier (Node). The dimensions and density of the channel conductances for these compartments are summarized in Table 1 and are partly based on previous EM studies (De Zeeuw *et al.*, 1990). Blocking of the different nodes was implemented by removing all active conductances at that node (Figure 5). The axial resistance was 150 Ωcm; the specific membrane capacitance (C_m_) was 1 µF/cm^2^ in all compartments except for the myelinated segments, where the C_m_ was 0.01 µF/cm^2^. The specific membrane resistivity (R_m_) was 12,000 Ωcm^2^ in all compartments except for the myelinated segments, where R_m_ was 1,200,000 Ωcm^2^ so that the myelin membrane time constant was 12 ms. When the myelin C_m_ was modified, *R*_*m*_ was set to a value that kept the membrane time constant at 12 ms.

**Table 1.** Two types of subthreshold

|  | Soma | Dendrites | Axon<br>Hillock<br>(AH) | Axon Initial<br>segment<br>(AIS) | Myelin | Node of<br>Ranvier |
| --- | --- | --- | --- | --- | --- | --- |
| Diameter ($\mu\text{m}$) | 20 | 1.5 | 2.7 - 1.3 | 1.3 | 1.3 | 1.3 |
| Length ($\mu\text{m}$) | 20 | 200 | 18 | 30 | 40 | 2 |
| $g_{\text{Na}}$ pS/$\mu\text{m}^2$ | 175 | 0 | 175 | 5000 | 0 | 5000 |
| $g_{\text{K}}$ pS/$\mu\text{m}^2$ | 50.638 | 0 | 50.638 | 300 | 0 | 300 |

### Model of axon with spines

Four spines were added to the axon model in the AH, 11.7, 15.3, 4.5 and 8.1 µm from the soma. Each spine had a neck length of 8.5, 4.5, 7, 3.5 µm, respectively, and a diameter of 0.2 µm. The spine head diameter and length were 2 µm each. The sodium and potassium conductances (gNa and gK, respectively) in the AH and in the spine heads were set as in the AIS (Table 1). For these simulations, the active conductances were removed from the Nodes of Ranvier.

### Building the IO network connectivity matrices

We constructed a network of IO composed of 1134 neurons randomly distributed within a volume of 250×500×200 µm, which resulted in 0.045 neurons per 10 µm^3^. We then clustered the neurons by their location using k-means clustering, and varied the number of neurons in a cluster by choosing k to be 1134 divided by the number of neurons in a cluster. The probability of a connection between two neurons decays with distance according to a Gaussian profile:

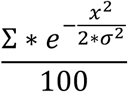

where ∑ is the maximal probability for connection (when the distance between the neurons is 0, x is the distance between neurons and σ sets the decay of connection probability with distance (see Figure 8H inset for examples). Note that the shape profile of neuron connectivity within a cluster could have a different ∑ and σ than the connectivity profile of neurons belonging to different clusters. The common neighbor distribution (Figure 8F, I) was calculated on randomly selected pairs of neurons within a distance of 40 µm.

### Constructing the IO network model

To simulate a realistic network of IO neurons, we followed the steps described above but with a few modifications. The network volume was 125×250×100 µm, and populated with 180 neurons (0.057 neurons for 10 µm^3^). These neurons were cloned from the three 3D-reconstructed olivary neurons, one of which is shown in Figure 5. We set the cluster size, k, to 40. ∑ and σ were 77 and 15 within a cluster and 45 and 20 between clusters, respectively (Figure 8H, inset, gray). The electrical connection between two neurons was mediated by 2 gap junctions (GJs). AGJ conductance (GJc) of 0.3 nS resulted in a coupling coefficient of 0.03±0.019 as in the experimental range (0.039±0.029). After adding GJc to the modeled cell, R_m_ was modified to maintain the experimental value of R_in_ (See details in (Amsalem *et al.*, 2016)). The spikes in the networks were created by current injection to the soma (simulated spikes) following a Poisson process.

## Supplementary Figures

**Figure S1.**
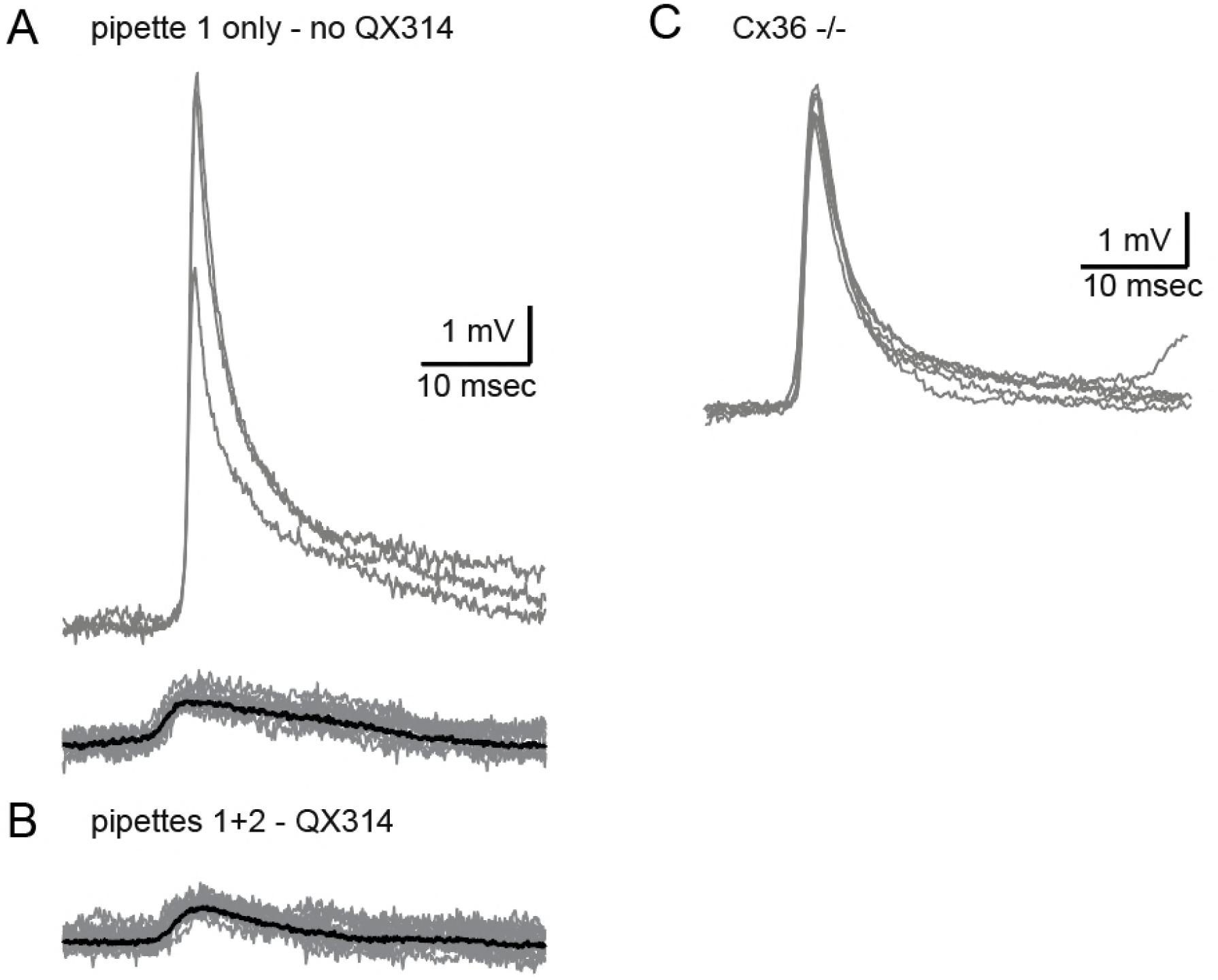
Related to Figure 4. The fast events are Internally Regenerative Events that are independent of gap junctions. (A) IREs (upper traces) and spikelets (lower traces) recorded from an olivary neuron with a regular internal solution. (B) breaking into the cell with another pipette, filled with intracellular solution + 2 mM QX 314, shows that with QX 314, IREs could not be detected while the spikelets’ shape and frequency were unaffected. (C) IREs recorded in CX 36 knockout mice showing that IREs can be generated in the absence of gap junctions.

**Figure S2.**
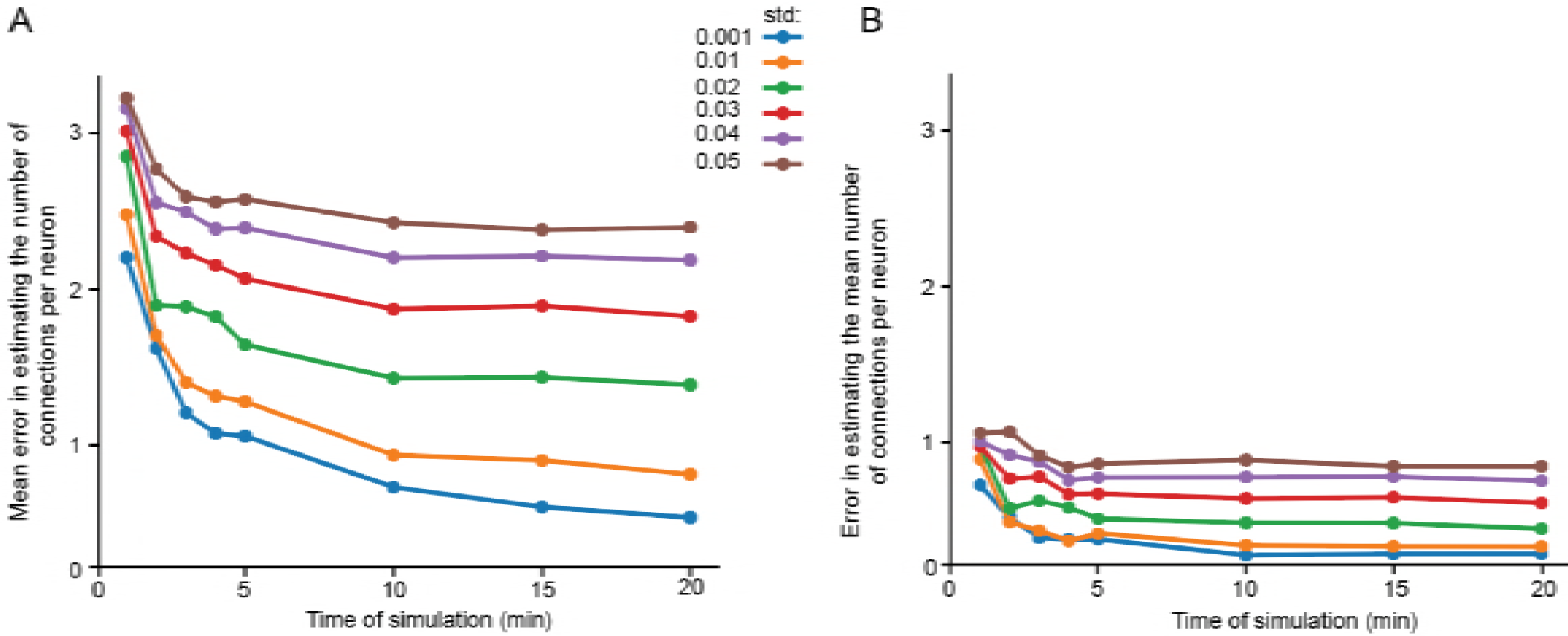
Related to Figure 8. The accuracy in estimating the number of connected neurons is strongly dependent on the variability in firing rate and the simulation duration. (A) difference between the estimated and real number of connections per neurons as a function of simulation duration for 6 different firing rate variabilities (std), color-coded as in the legend. (B) error in estimating the mean number of connections in the network (plus sign in Figure 8C) as a function of simulation duration for 6 different firing rate variabilities (std). The calculation was done only on neurons that had common neighbors (n=278 pairs). Mean firing rate was 0.058 Hz.

**Figure S3.**
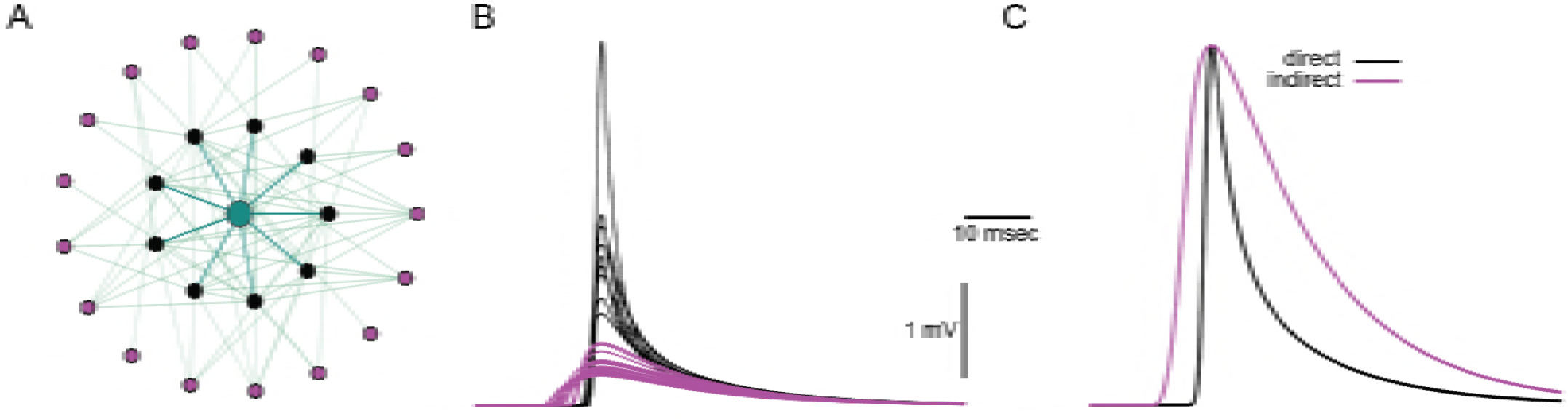
Related to Figure 5 and Figure 6. Difference in events waveform can be attributed to spikes form directly and indirectly connected cells. (A) modelled IO network in which a green cell was connected directly to 9 other cells (black circles), and indirectly to 17 other cells (purple circle). (B) spikelets in this cell due to spikes in directly connected cells (black traces) or indirectly connected cells (purple traces). (C) average normalized spikelets from B. This network is similar to the network in Figure 8A, with two modification - the number of GJs was changed to 6 and the GJc to 0.25 nS.

